# sEEG-Suite: An Interactive Pipeline for Semi-Automated Contact Localization and Anatomical Labeling with Brainstorm

**DOI:** 10.1101/2025.09.16.676602

**Authors:** Chinmay Chinara, Raymundo Cassani, Takfarinas Medani, Anand A. Joshi, Samuel Medina Villalon, Yash S. Vakilna, Johnson P. Hampson, Kenneth Taylor, Francois Tadel, Dileep Nair, Christian G. Bénar, Sylvain Baillet, John C. Mosher, Richard M. Leahy

## Abstract

Stereoelectroencephalography (sEEG) is a critical tool for mapping epileptic networks in patients with drug-resistant epilepsy. Accurate localization and labeling of sEEG contacts are essential for identifying the seizure onset zone (SOZ) and ensuring optimal resective surgery. Traditional methods for localizing and labeling sEEG contacts rely on manual processing, which is prone to human error and variability. To address these challenges, we developed and integrated a semi-automatic sEEG contact localization and labeling pipeline within Brainstorm^1^, an open-source software platform for multimodal brain imaging analysis^2–8^, widely adopted in the neuroscience community with [over 50,000 registered users](https://neuroimage.usc.edu/brainstorm/Community) and an active online [user forum](https://neuroimage.usc.edu/forums/). The software has been [supported](https://neuroimage.usc.edu/brainstorm/Introduction#Support) by the National Institute of Health (NIH) for over two decades. The pipeline presented in this paper performs three key steps: (1) import and apply rigid co-registration of post-implantation Computed Tomography (CT) or post-CT with pre-implantation Magnetic Resonance Imaging (pre-MRI), (2) post-CT image segmentation and semi-automatic detection of sEEG contacts using GARDEL^9^, which has been integrated as a Brainstorm plugin, and (3) automatic anatomical labeling of contacts using standard and commonly used brain anatomy templates and atlases. Integrating this pipeline into Brainstorm brings the best of both worlds: GARDEL’s automation and Brainstorm’s multimodal data compatibility, and rich library of visualization and advanced analysis tools at both sensor and source level^6^. This sEEG-Suite tool facilitates reproducible research, supports clinical workflows, and accelerates sEEG-based investigations of invasive brain recordings.

## Tutorials and Documentation

Brainstorm offers a comprehensive collection of [detailed tutorials](https://neuroimage.usc.edu/brainstorm/Tutorials) that cover all major components of the platform, supporting users from basic data processing to advanced multimodal analyses. Within this ecosystem, the sEEG-suite is accompanied by its own dedicated tutorials, with each section of the suite linking to a corresponding resource. For convenience, the complete set of tutorials for stereo-electroencephalography (sEEG) analysis can be accessed here: sEEG-suite: Stereo-electroencephalography (sEEG) Analysis.

Figure 1 illustrates the complete end-to-end workflow for sEEG analysis in Brainstorm. The process begins with the acquisition of anatomical and implantation imaging, including both pre- and post-MRI and post-CT scans. The CT image is rigidly co-registered to the MRI to ensure alignment of anatomical and electrode information, after which skull stripping is applied to remove non-brain structures and CT artifacts. From the CT, an intensity thresholded isosurface is generated to identify candidate electrode contacts that will be used during the localization process. At this stage, Brainstorm offers two localization strategies: a manual interface that allows users to add/remove/merge/rename electrodes and add/remove contacts, and an automated localization procedure using GARDEL, which streamlines contact detection and reduces the need for extensive manual intervention. Once preliminary electrode positions are obtained, users can refine and correct them to generate a finalized set of electrode coordinates. These coordinates are then subjected to automated anatomical labeling, in which each contact is mapped to underlying cortical and subcortical structures based on atlas information derived from the MRI segmentation or provided from the different atlases available in Brainstorm. The localized and labeled electrode contacts can further be linked to the raw electrophysiological recordings, enabling Brainstorm to integrate anatomy with the corresponding time series data.

**Figure 1.**
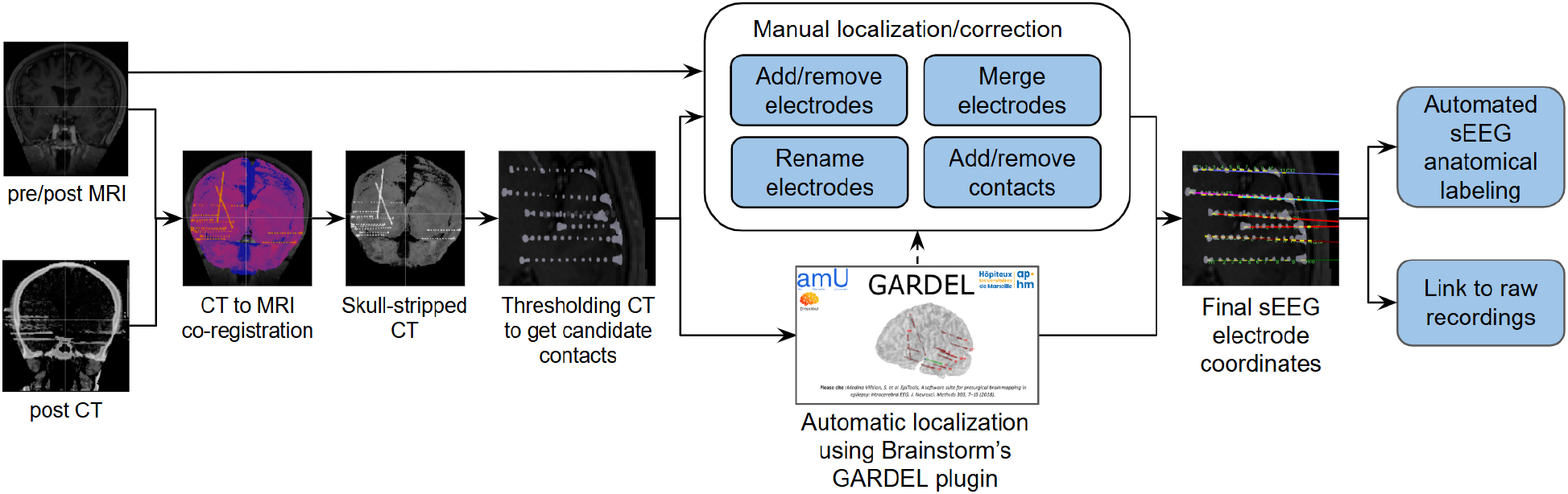
End-to-end flowchart for sEEG analysis in Brainstorm

## Anatomy and preprocessing

pre-MRI (Figure 2a) and post-CT (Figure 2b) scans are acquired for each subject. The post-CT volume is rigidly co-registered with the pre-MRI volume using the Statistical Parametric Mapping (SPM) or USC’s CT2MRI (available as a Brainstorm plugin) (Figure 2c). The pre-MRI serves as an anatomically grounded reference for subsequent processing. Specifically, the pre-MRI is used to perform skull-stripping of the post-CT using tissue segmentation obtained either by BrainSuite^10^ or SPM^11^ (using function mri_skullstrip.m), effectively removing extracranial tissues and eliminating artifacts associated with the sEEG electrode wire bundles (Figure 2d). The resulting artifact-free post-CT volume is then subjected to an automatic intensity thresholding to detect and extract metal artifacts, which are potential candidates for electrode contact localization (Figure 2e). More details can be found in the tutorial: [CT to MRI co-registration](https://neuroimage.usc.edu/brainstorm/seeg/ct2mri).

**Figure 2.**
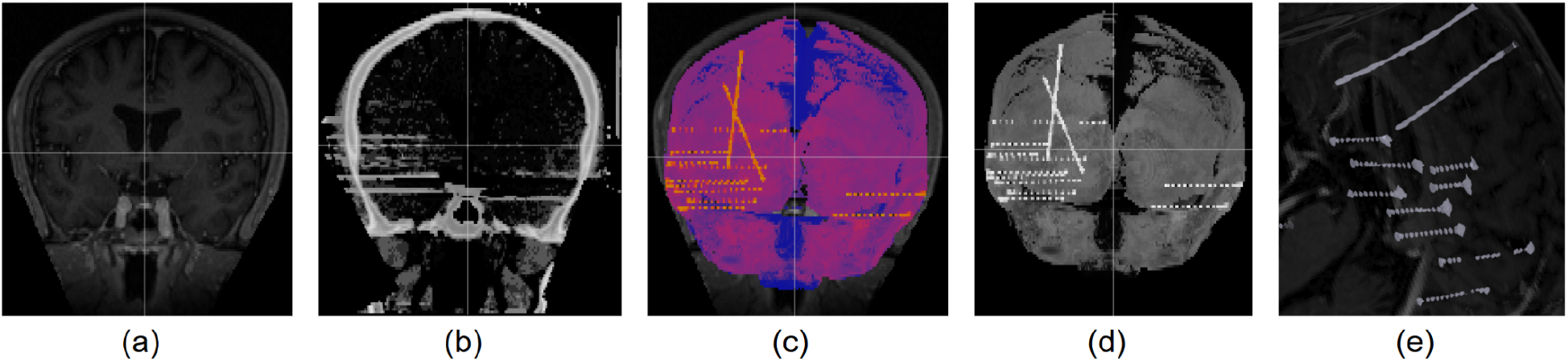
(a) pre-MRI; (b) post-CT; (c) post-CT co-registered to pre-MRI and skull-stripped using SPM (the colored image is the post-CT overlaid on the pre-MRI to show their proper alignment); (d) Skull-stripped post-CT by itself; (e) Isosurface generated from skull-stripped post-CT using intensity thresholding to capture metal artifacts as candidate contacts

## Manual contact localization

Prior to localizing the electrode contacts, knowledge of the implantation scheme is required in order to have the correct naming convention (used by the center/neurosurgeons) for the various electrodes used. Brainstorm’s iEEG graphical panel allows creating an electrode implantation manually by first defining the electrode model and constructor (through a list of predefined uniformly spaced electrode models from various constructors, such as [DIXI, PMT](https://diximedus.com/), etc, or users can choose to define and load their own model). The user can then set the electrode tip and skull entry to render the electrode (Figure 3). This process can be done in the 3D space (using the intensity thresholded mesh generated in the previous section), in the 2D MRI viewer (using the intensity thresholded post-CT overlaid on the pre-MRI), or only on post-MRI if the CT is not available. More details can be found in the tutorial: [SEEG contact localization and labeling](https://neuroimage.usc.edu/brainstorm/seeg/SeegContactLocalization).

**Figure 3.**
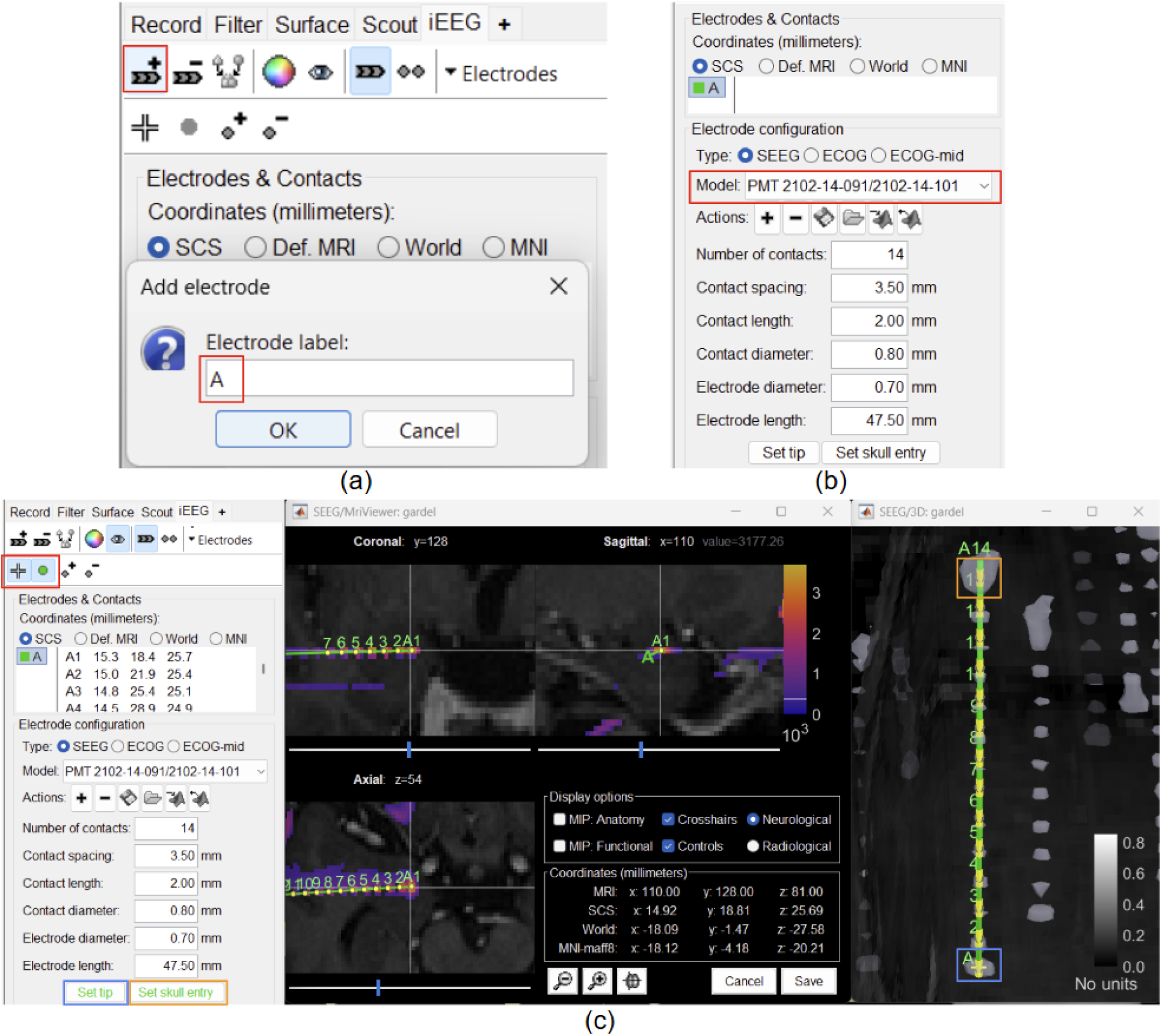
(a) Create an electrode and assign label to it (in red); (b) Define the electrode model (in red); (c) On the 3D figure (SEEG/3D: gardel), set the electrode tip (in blue) and skull entry (in orange) using the surface selection button (in red) to render the electrode both in 2D MRI viewer and 3D figure

## Automatic contact localization

With the click of a single button inside Brainstorm (Figure 4), we have live, interactive visualization of the detection process, enabling users to observe contact identification, grouping, and ordering in real time. The electrode names are automatically assigned from **A-Z, AA-ZZ**, etc., in the order they are detected, which can be renamed as desired during post-processing (Figure 5). On the button click, GARDEL uses the skull-stripped post-CT volume along with the intensity threshold to identify high-density regions corresponding to metallic artifacts that are considered as electrode contacts. These detected points are grouped into individual leads, with contacts sorted along each electrode trajectory, designating the deepest contact as the first in sequence. This integration reduces the need for manual contact detection and electrode indexing, significantly streamlining the localization workflow. More details can be found in the tutorial: [Automatic SEEG Contact Localization using GARDEL](https://neuroimage.usc.edu/brainstorm/Tutorials/AutoContactLocGardel).

**Figure 4.**
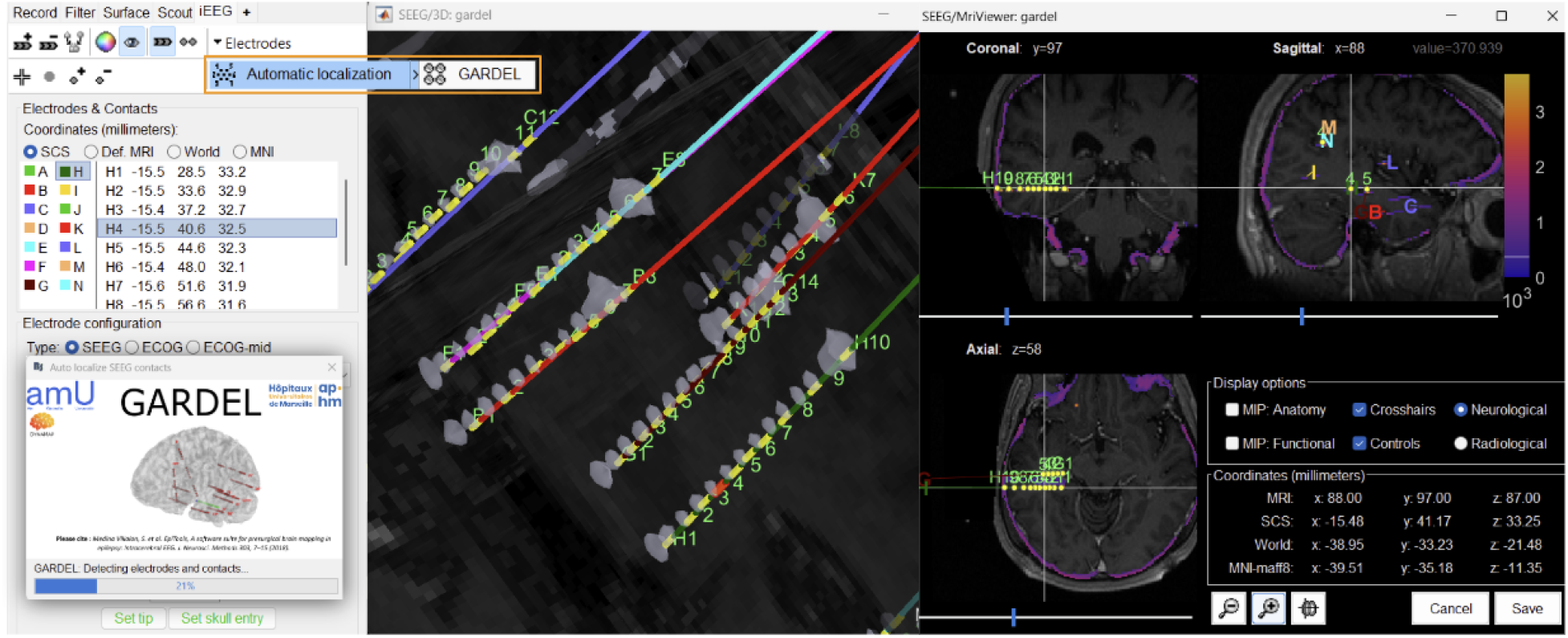
The Brainstorm interface displaying the automatically detected contacts, with the GARDEL button (in orange) that triggers the automatic detection

**Figure 5.**
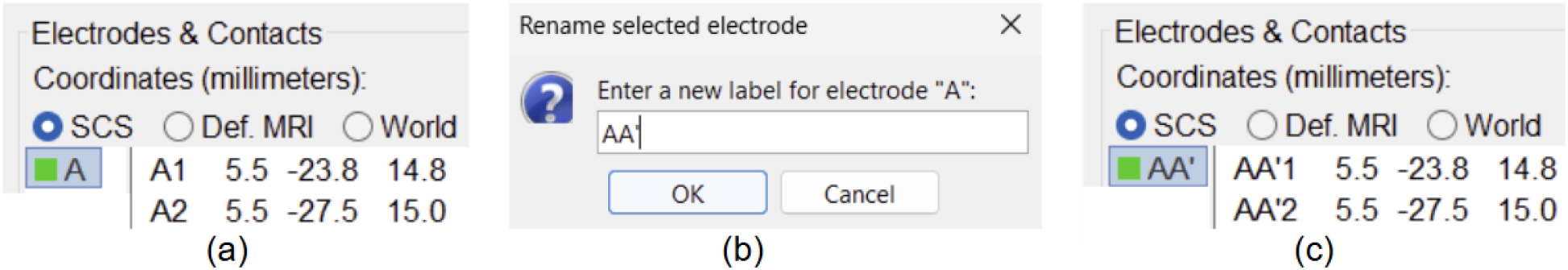
Renaming electrode (a) Electrode **A** before renaming; (b) Brainstorm interface to rename the electrode (double click on electrode **A** and change name to **AA’**); (c) Electrode A renamed to **AA’** (along with all the contacts)

## Post-processing of the electrodes/contacts

All the necessary post-processing can be done using the Brainstorm interface interactively. If there is a wrongly detected electrode, we can delete and manually rectify it by creating it from scratch (as mentioned in the Manual contact localization section above) along with renaming them if needed (Figure 5). There could be cases where a single electrode may be detected as multiple in which case we need to merge them as one (Figure 6). We also allow fine-tuning at the contact level, where we can add/remove them (Figure 7-8). On doing any of the editing operations, it is also ensured that the ordering of the contacts is maintained with the deepest contact as first in the sequence. More details can be found in the tutorial section: [Edit the contacts positions](https://neuroimage.usc.edu/brainstorm/seeg/SeegContactLocalization#Edit_the_contacts_positions).

**Figure 6.**
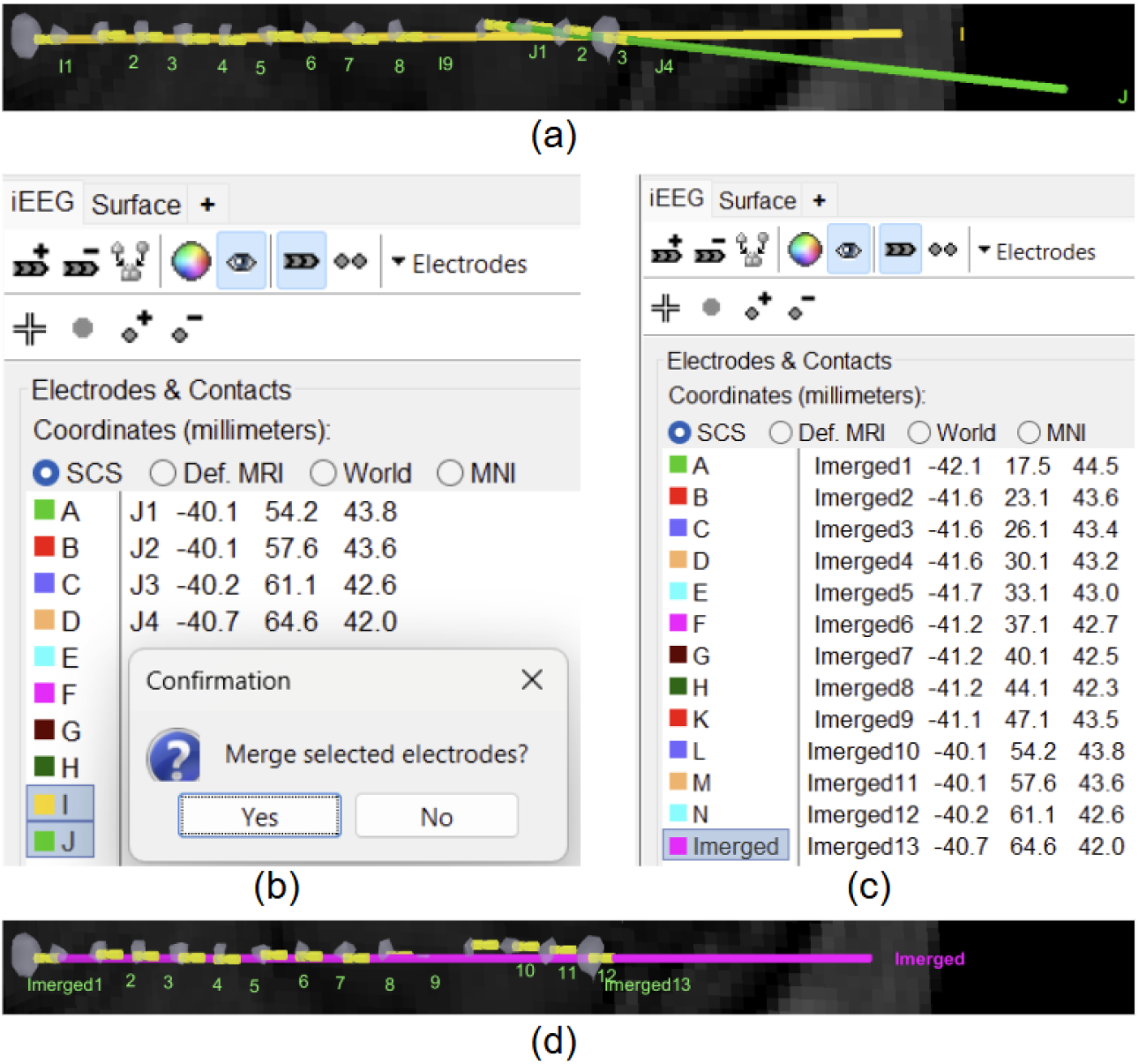
Merging electrodes using the Brainstorm iEEG panel; (a) Electrodes **I** and **J** are wrongly detected as separate electrodes; (b) Brainstorm interface showing option for merging them; (c) The merged electrode in iEEG panel (**I** and **J** get replaced by **Imerged**); (d) The rendered merged electrode

**Figure 7.**
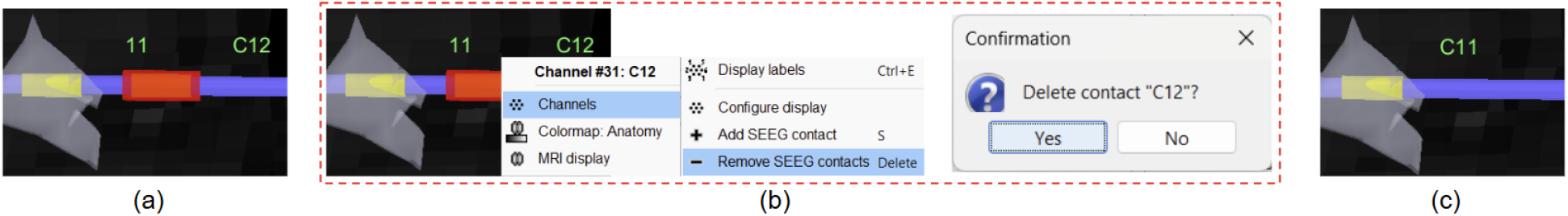
Removing a contact using 3D figure; (a) Wrongly detected contact (selected contact **C12** in red); (b) Brainstorm interface showing option for removing it (right click on the **C12** contact to get this menu); (c) Contact removed

**Figure 8.**
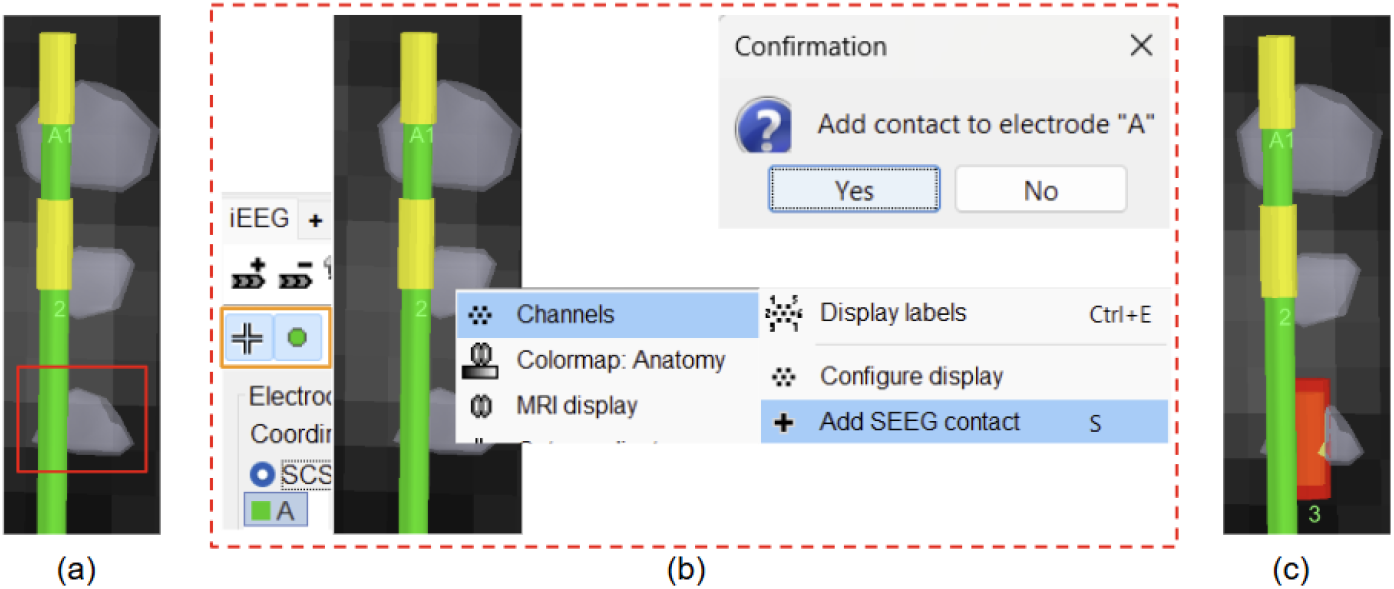
Adding a contact using 3D figure; (a) Missed detecting contact (3rd from top for electrode **A** in red); (b) Brainstorm interface showing option for adding it (turn on surface/centroid selection and select electrode **A** in iEEG panel, select the surface point and right click on it to get this menu); (c) Missing contact **A3** added

## Automated anatomical labeling

Brainstorm supports multiple standardized brain anatomy templates (ICBM152, Colin27, USCBrain, FsAverage, etc.) and [brain atlases](https://neuroimage.usc.edu/brainstorm/Tutorials/DefaultAnatomy) (AAL, Desikan-Killiany, Brainetome, Schaefer, etc.), all of which can be added from the interface as shown in Figure 9(a-b). The spatial coordinates of contacts localized by the steps above are automatically cross-referenced against these atlases, resulting in detailed anatomical labels (e.g., specific cortical gyri, subcortical nuclei) as shown in Figure 9(c-d). This enriched labeling framework provides a comprehensive anatomical context for each contact (the region of the brain the contact belongs to), facilitating both clinical interpretation and research analyses. More details can be found in the tutorial section on [anatomical labeling](https://neuroimage.usc.edu/brainstorm/Tutorials/Epileptogenicity#Anatomical_labelling). Work by^12^ demonstrates a clinical validation study using this feature of Brainstorm.

**Figure 9.**
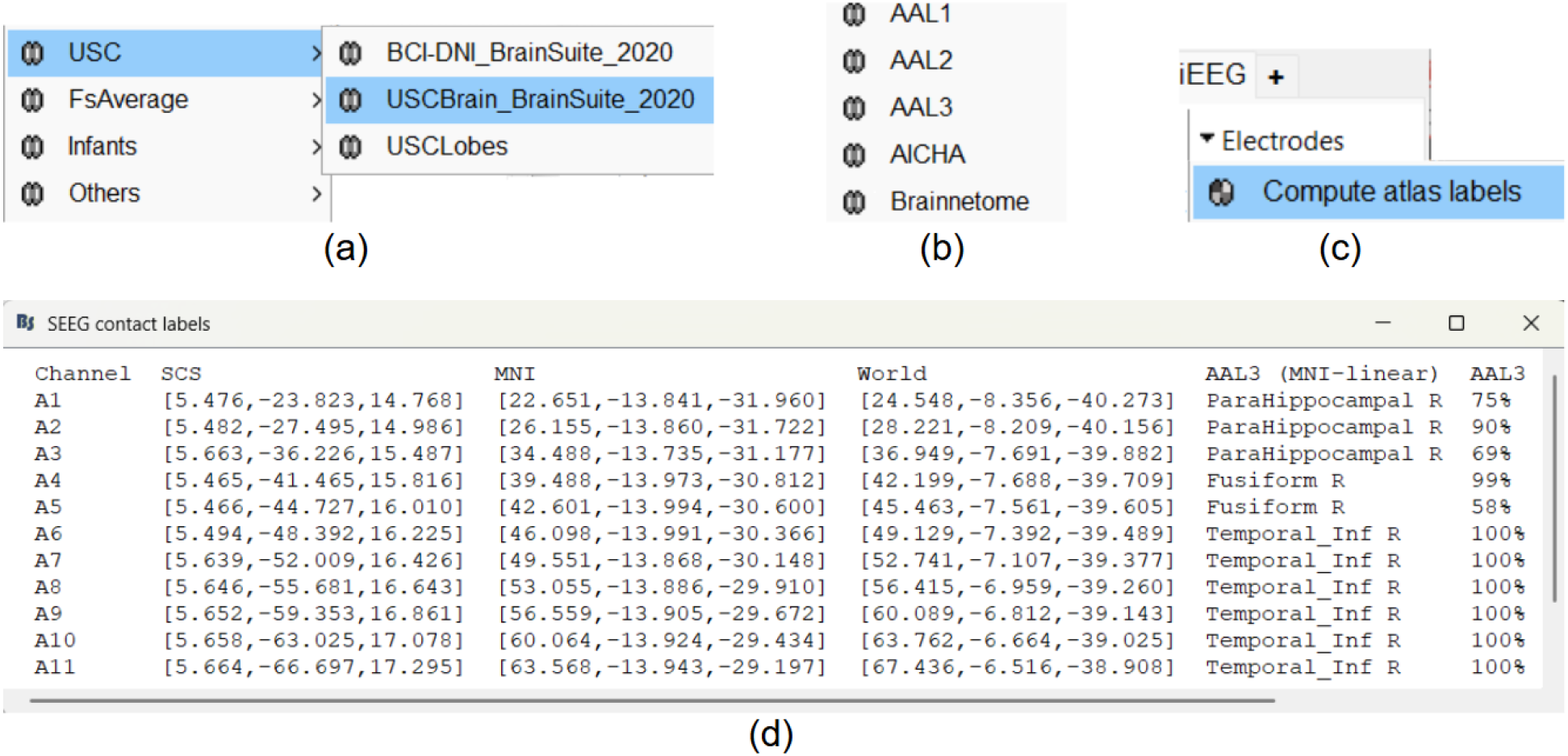
(a) The different brain anatomy templates supported inside Brainstorm (b) The different brain atlases supported inside Brainstorm (c) Brainstorm interface to compute the anatomical labeling (in the iEEG panel select the electrode **A**, go to *Electrodes > Compute atlas labels*) (d) The computed anatomical labels for contacts in electrode **A** using the AAL3 atlas

## Link to raw recordings

Once the sEEG contacts have been localized, we can associate them with the corresponding raw electrophysiological recordings. This link ensures that anatomical localization can be directly tied to the recorded brain signals. To establish this connection, it is essential that the contact names match exactly with the channel names in the raw recordings. Any mismatch (such as differences in naming conventions, capitalization, or numbering) will prevent Brainstorm from aligning the contacts with the appropriate channels. Detailed instructions on how to perform this linking can be found in the tutorial section: [Access the recordings](https://neuroimage.usc.edu/brainstorm/seeg/SeizureFingerprinting#Access_the_recordings). Following these guidelines ensures that each localized contact is correctly mapped to its electrophysiological data, allowing for reliable analysis and visualization. Once this step is completed, users can proceed either with the sEEG data analysis within Brainstorm using its [rich and complete set of tools](https://neuroimage.usc.edu/brainstorm/Tutorials#sEEG-suite:_Stereo-electroencephalography_.28sEEG.29_analysis) or export the final data into any of the [supported formats](https://neuroimage.usc.edu/brainstorm/Introduction#Supported_file_formats) for analysis with other software.

## Statement of Need

Intracranial electrode localization, particularly for sEEG, is a foundational step in epilepsy surgery planning and neuroscience research. At present, many researchers and clinicians rely on manual workflows or a patchwork of separate tools for localization, which results in time-consuming procedures, inter-operator variability, and fragmented pipelines. Manual identification and labeling of electrode contacts can take several hours per subject, slowing down the workflows, while subjective differences between operators introduce inconsistencies in results. Moreover, existing practices often require multiple platforms, i.e., one for CT-MRI co-registration, another for electrode detection, and another for anatomical labeling, and another for data analysis, thereby adding unnecessary complexity and risk of errors in the final results.

Integrating GARDEL with Brainstorm provides seamless, one-click deployment of intracranial electrode localization directly within Brainstorm, eliminating the need for external scripts and reducing manual intervention. This reduces localization time from hours to minutes (with some minor post-processing), enabling researchers to efficiently scale analyses across larger patient cohorts. The integration also enhances reproducibility by standardizing detection parameters, minimizing manual intervention, and unifying the entire workflow of localization and labeling with the advanced analysis of electrophysiological data into a single interactive environment.

While other tools, such as DEETO^13^, FASCILE^14^, LeGUI^15^ and 3D Slicer’s sEEG Assistant^16^ have demonstrated dramatic speedups compared to manual localization, they do not provide an end-to-end processing workflow from anatomical to functional analyses like Brainstorm that does.

## Acknowledgements

Research reported in this publication was supported by the National Institute of Biomedical Imaging and Bioengineering (NIBIB) of the National Institutes of Health (NIH) under award numbers R01EB026299 and RF1NS133972, DOD award HT94252310149.

